# Caspofungin binding to iron compromises its antifungal efficacy against *Candida albicans*

**DOI:** 10.1101/2024.12.30.630750

**Authors:** Andreia Pedras, Cláudia Malta Luís, Luís M. P. Lima, Dalila Mil-Homens, Catarina Amaral, Américo G. Duarte, Wilson Antunes, Ana Gaspar-Cordeiro, Ricardo O. Louro, Pedro Lamosa, Cláudio M. Soares, Diana Lousa, Catarina Pimentel

## Abstract

Echinocandin drugs, such as caspofungin, inhibit the synthesis of β-1,3-D-glucans, which are essential components of the fungal cell wall. These drugs are often the preferred option for treating invasive fungal infections (IFIs) caused by *Candida* spp. due to their superior efficacy compared to other antifungal agents. Iron overload conditions, which exacerbate fungal burden, are well-documented as significant risk factors for the progression of IFIs. Recent *in vitro* studies have suggested that iron overload may also reduce the efficacy of cell wall-perturbing agents, such as echinocandins, against *Candida albicans*, by altering the composition of the fungal cell wall. Here, we show that iron loading conditions which do not interfere with the cell wall composition are still capable of recapitulating the caspofungin-resistant phenotype induced by iron in *C. albicans*. Spectroscopic analyses provided evidence that caspofungin binds to iron through its ethylenediamine moiety and two amide groups. Consistent with the *in vitro* activity of β-1,3-D-glucan synthase, molecular dynamics simulations revealed that, when bound to iron, caspofungin undergoes conformational changes that may reduce its ability to inhibit the enzyme.

Importantly, the *in vivo* antifungal efficacy of caspofungin is compromised in a *Galleria mellonella* model of IFI caused by *C. albicans* simulating a context of iron overload. This effect may extend beyond *C. albicans* infections, as the antagonism between iron and caspofungin was also observed in other medically important fungi causing IFIs.

## 1. Introduction

Invasive fungal infections (IFIs), in which fungi enter the bloodstream and/or invade internal organs [1], are responsible for approximately 1.5 million deaths worldwide each year [2]. Among them, those caused by the yeast *Candida*, known as invasive candidiasis, are the most prevalent in hospitalized patients undergoing immunosuppressive treatments or intensive antibacterial therapies [3, 4].

*Candida* spp. colonize multiple tissues in the human body but can switch from commensals to pathogens under specific conditions, driven by several virulence factors. One key factor is their ability to form robust biofilms [5], which enable them to adhere and proliferate on host tissues and medical devices [6]. Bloodstream infections caused by *Candida* spp., referred to as candidemias, are common manifestations of invasive candidiasis and present a serious challenge in Intensive Care Units (ICUs), where two thirds of cases occur [4, 7]. These infections are associated with high rates of mortality and morbidity and often lead to prolonged hospital stays, placing a significant burden on individuals and healthcare systems [8]. Among patients with candidemia, *Candida albicans* is the most frequently identified species, followed by *Candida glabrata* and *Candida parapsilosis* [9].

Several antifungal drugs are available to treat invasive candidiasis [10]. Azoles are the most commonly used antifungal class due to their minimal side effects, low cost and broad spectrum of activity [10, 11]. Azole drugs act by inhibiting the biosynthesis of ergosterol, an important component of fungal membranes, thereby arresting cell growth and division [11–13]. However, several factors limit their widespread use: some *Candida* spp. are inherently resistant to these drugs, their intensive use has led to the emergence of resistant clinical isolates, and their fungistatic rather than fungicidal nature restricts their efficacy, particularly in immunocompromised patients [14, 15]. As a consequence, the echinocandin class of antifungals has increasingly become the first-line treatment for invasive candidiasis in many cases [10]. Echinocandin drugs, such as caspofungin, micafungin and anidulafungin, are fungicidal lipopeptide molecules that inhibit the catalytic Fks subunit of the β-1,3-D-glucan synthase, the enzyme responsible for synthesizing β-1,3-D-glucans, a critical component of the inner layer of the fungal cell wall [16].

The fungal cell wall plays a crucial role in immune recognition, as its components function as pathogen-associated molecular patterns (PAMPs) that are recognized by pattern recognition receptors (PRRs) on host cells [17]. It is therefore not surprising that antifungal agents disrupting cell wall architecture, such as echinocandins, also affect the interaction of *Candida* spp. with the host [18–22].

Iron availability is another key factor influencing the host-fungus interaction. While the host employs an iron-withholding strategy to restrict the growth of invading microorganisms [23], this iron-depleted environment triggers changes in the fungal cell wall that promote immune evasion [21, 24, 25]. As expected, iron overload is often linked to increased fungal burden and poor clinical outcomes [26–29]. These conditions are relatively common and can result from diseases such as hereditary hemochromatosis, hepatic disease, or cancer, as well as from medical treatments like chemotherapy or multiple blood transfusions [30].

Iron overload also decreases *C. albicans* susceptibility to agents targeting the cell wall [26, 31]. This effect has been attributed to iron-mediated alterations in key cell wall components, including chitin, mannans, and glucans, which have been proposed as the underlying mechanism for the increased resistance to those agents [26, 31]. In this study, under iron overload conditions that did not significantly alter cell wall composition, the same phenotype, was observed, suggesting the involvement of additional, still unknown mechanisms. While iron alone does not directly interfere with the *in vitro* activity of β-1,3-D-glucan synthase, it prevents the inhibition of the enzyme by caspofungin. Since caspofungin can bind iron, altering its conformation, we propose that this change impairs the inhibitory activity of caspofungin on β-1,3-D-glucan synthase. The potential biological significance of our work is supported by the observation that the *in vivo* antifungal efficacy of caspofungin is compromised in a *Galleria mellonella* model of invasive candidiasis when the larvae are preloaded with iron to mimic iron overload disorders.

## 2. Materials and Methods

### Strains and growth conditions

The strains used in this study are listed in Table S1. Yeast species were maintained on YPD agar plates. Unless otherwise stated, all assays were performed in SC citrate medium at pH 6.5 (Complete Supplement Mixture: 0.77 g/L; yeast nitrogen base w/o amino acids and ammonium sulphate: 1.7 g/L; ammonium sulphate: 5.4 g/L; glucose: 2%; citrate buffer: 5 mM). For all assays, unless otherwise specified, *C. albicans* SC5314 cultures were grown to exponential phase (OD_600_ 0.8) and treated with 5 mM FeSO_4_, 0.375 µg/mL caspofungin or both, for 24 h at 30 °C. *Aspergillus fumigatus* was grown at 37 °C on minimal medium with 1% glucose and 20 mM glutamine as nitrogen source. Conidia were harvested and kept in saline buffer (0.01% tween-80 and 0.9% NaCl). For spot dilution assays, 5 µL of serial dilutions (ranging from 8 × 10^6^ to 80 cells/mL) of exponentially growing cultures were spotted onto SC agar plates supplemented with indicated compounds and the plates were incubated at 30 °C for 48 h. For spot dilution assays with iron-pretreated cultures, precultures and cultures were grown in medium supplemented with the specified concentrations of FeSO₄ and washed with PBS before spotting. For growth curves, *C. albicans* SC5314 (0.01 OD_600_) were supplemented with the indicated compounds, incubated at 30 °C for 65 h (BioTek Synergy Neo2) and OD_600_ was measured every 1 h. Assays were performed using biological triplicates and technical quadruplicates.

### Cell viability assay

*C. albicans* SC5314 cells, cultured as described above, were harvested by centrifugation, washed with PBS, and resuspended in 50 μg/mL propidium iodide (PI). The assays were performed using the S3e cell sorter (Bio-Rad), with the PI signal detected using the 488 nm excitation laser and the FL3 detection channel. Four biological replicates were analyzed for each condition, with a total of 10^5^ cells counted per sample.

### Scanning electron microscopy

*C. albicans* SC5314 cultures, grown as indicated above, were harvested by centrifugation and fixed in a solution of 0.4% glutaraldehyde and 4% formaldehyde in sodium cacodylate buffer 0.1 M for 18 h at room temperature. Following three washing steps with sodium cacodylate buffer, the samples were sequentially dehydrated in ethanol solutions of increasing concentrations (50%, 70%, 90% and 100%). Ethanol was replaced with tert-butyl alcohol and the samples were incubated at 30 °C for 1 h, then placed on ice until fully frozen, lyophilized and stored at room temperature until the SEM analysis. Samples were gold sputtered with an 8 nm layer using an electron sputter (Cressington 108) and imaged with a Hitachi SU8010 scanning electron microscope, at 1.5 kV.

### Checkerboard assays

The assays were performed in SC citrate medium using 96-well flat bottom plates, according to the CLSI standard methods M27-A3 and M38 for yeast and filamentous fungi, respectively [32, 33], as described in [34, 35], with minor modifications. Stock solutions of caspofungin and micafungin were prepared in ultrapure water and the stock solution of anidulafungin was prepared in DMSO. FeSO_4_ stock solution was freshly prepared in bi-distilled water and filtered. Two-fold serial dilutions were prepared for each compound, resulting in final concentration ranges from 1.5 to 0.0015 µg/mL for caspofungin, micafungin and anidulafungin, and 50 to 0.78 mM for FeSO_4_. For yeast, a 2 × 10^6^ cells/mL suspension obtained from a single colony was diluted to 3 × 10^3^ cells/mL (6 × 10^3^ cells/mL for *S. cerevisiae*) in medium, to prepare the final inoculum. For *A. fumigatus*, a 10^5^ spores/mL suspension was used, and 10^4^ spores were placed in each well. Growth was recorded after 24 and 48 h at 30 °C (or 37 °C for *C. glabrata* and *A. fumigatus*), by measuring OD_600_ (yeast strains) or using phase contrast microscopy (*A. fumigatus*). For the latter, a Wild Heerbrugg M7A stereo zoom microscope was used and images were captured using a digital camera. The minimum inhibitory concentration (MIC) endpoints were defined as the lowest drug concentration resulting in a growth reduction of more than 50%, for yeast strains. The minimum effective concentration (MEC) was defined as the lowest concentration leading to a visible reduction in spore germination for *A. fumigatus*. The type of interaction for each metal/drug combination was determined by calculating the fractional inhibitory concentration (FIC) index (ƩFIC = FIC_Fe_ + FIC_drug_). A ƩFIC > 4 indicates an antagonistic effect. For each metal/drug combination, the FIC was calculated as follows: FIC_Fe_ = MIC_Fe+drug_/MIC_Fe_ and FIC_drug_ = MIC_drug+Fe_/MIC_drug_. Checkerboard assays were repeated at least twice.

### Biofilm assays

Biofilm formation was monitored using the protocol described in [36] with minor modifications. *C. albicans* SC5314 cultures were grown overnight in SC citrate medium at 30 °C with agitation. Prior to the assays, the wells of surface treated 96-well flat bottom plates were conditioned with 50% fetal bovine serum for at least 30 min [37]. After washing with PBS, *C. albicans* cells were added at a final OD_600_ of 0.5 in 200 µL of SC citrate medium and the plates incubated at 37 °C with agitation for 90 min (adherence step). Non-adherent cells were aspirated, the wells washed with PBS, and following the addition of 200 µL of fresh medium, the plates were incubated at 37 °C with agitation for 24 h (growth step). Caspofungin and FeSO_4_ or FeCl_3_ were added at the beginning of the adherence step (sustained inhibition assay), after the adherence step (developmental inhibition assay), or at the end of the growth step (mature biofilm disruption assay). Following the growth step, the medium was aspirated, biofilms were washed with PBS, and biofilm viability assays were performed as described in [37]. Briefly, 100 µL of 0.5 mg/mL 3-(4,5-dimethyl-2-thiazolyl)-2,5-diphenyl-2H-tetrazolium bromide (MTT) in PBS were added to each well and the plates were incubated at 37 °C for 4 h. Crystals were solubilized in 100 µL of acid isopropanol and incubated at room temperature on a shaker, for 15 min and OD_550_ was measured. The sessile minimum inhibitory concentration (SMIC) was defined as the lowest drug concentration resulting in a biofilm viability reduction of more than 50%. All assays were performed with biological quadruplicates and technical triplicates.

### Confocal microscopy

*C. albicans* SC5314 cultures, grown as described above, were harvested by centrifugation, washed and resuspended in PBS. Fluorescent labelling of yeast cell wall components was done as described in [38]. For chitin staining, cells were incubated with calcofluor white (CFW) at a final concentration of 5 µg/mL, for 10 min. For β-1,3-D-glucan staining, cells were incubated with aniline blue (AB) at a final concentration of 1 mg/mL, for 30 min. For mannans, cells were stained with concanavalin A (ConA) at a final concentration of 100 µg/mL, for 30 min. After incubation, cells were washed and resuspended in PBS. Images were acquired using a Zeiss LSM 880 point scanning confocal microscope controlled by Zeiss Zen 2.3 (black edition) software.

### Quantification of cell wall components

Quantification of β-1,3-D-glucans was performed as described in [39] with modifications. *C. albicans* SC5314 cells were grown as indicated above, harvested, washed with TE buffer (10 mM Tris-HCl, 1 mM EDTA, pH 8.0), and dried at 80 °C for 24 h. Samples were resuspended in TE buffer supplemented with 1 M NaOH, incubated at 80 °C for 30 min, an aniline blue solution (0.03% aniline blue, 0.18 M HCl, 0.49 M glycine-NaOH, pH 9.5) was added. After incubation at 50 °C for 30 min, samples were cooled to room temperature for 30 min, centrifuged, and fluorescence of the supernatant was measured in a white opaque 96-well microplate using a fluorescence plate reader (FLUOstar OPTIMA; BMG LABTECH), with an excitation wavelength of 400 nm and an emission wavelength of 460 nm. For chitin quantification, cells were stained with CFW and analyzed by confocal microscopy, as described above. Fluorescence intensities of stained cells (n ≥ 100) were quantified using Fiji [40]. Assays were performed in biological triplicates. For mannans, cells were stained ConA, as described above, filtered and analyzed by flow cytometry. Mannans were quantified using an S3e cell sorter (Bio-Rad), using the 488 nm excitation laser and the FL3 detection channel. Four biological replicates were analyzed per condition and a total of 10^5^ cells were counted per sample.

### Iron binding assays

Assays were performed using FeCl_3_ due to its chemical stability in aqueous solutions. Stock solutions of caspofungin acetate and micafungin sodium salts were prepared in water at 2 mg/mL and 5 mg/mL, respectively. A 500 mM solution of NaSCN (thiocyanate) was prepared in water. A working solution of FeCl_3_ (5 mM) was prepared by diluting a concentrated stock solution (25.6 mM) previously standardized via complexometric titration. The spectra of individual components were plotted in aqueous solutions as follows: caspofungin at 150 µM, micafungin at 30 µM, NaSCN at 7.5 mM and FeCl_3_ at 150 µM. Competitive binding assays between thiocyanate and caspofungin or micafungin were performed using samples containing 7.5 mM of NaSCN and 150 µM of FeCl_3_, by stepwise addition of caspofungin or micafungin up to 1.2 equivalents of the metal concentration. UV-Visible absorption spectra were acquired at 25 °C on a PerkinElmer lambda 650 UV-Visible spectrophotometer in 1 cm path length quartz cuvettes. A stock solution of 100 mM 3-(N-morpholino)-propanesulfonic acid (MOPS) buffer was prepared at pH 7 in water, and used in the UV spectrophotometric titrations performed in aqueous medium buffered at pH 7 with 20 mM MOPS. For the interaction between iron and caspofungin, a sample with 200 µM of FeCl_3_ was titrated by stepwise additions of caspofungin up to 2.0 equivalents of the metal concentration (15 titration points), over a wavelength range of 260–360 nm. For the interaction between iron and micafungin, a sample containing 30 µM of FeCl_3_ was titrated by stepwise additions of micafungin up to 2.6 equivalents of the metal concentration (13 titration points), over a wavelength range of 280–330 nm. UV-visible absorption spectra were acquired for each of the individual titration points, as described above. Conditional binding (complexation) constants (log *K*_f_) for each titration were determined using the HypSpec software [41, 42], which applies the combined UV titration data to calculate the constant and corresponding standard deviation for each interaction. In each titration, the UV spectra of sample solutions of iron and caspofungin or micafungin were used as standards for data fitting. A simple 1:1 binding stoichiometry was initially assumed for the interactions of both caspofungin and micafungin with iron, and this assumption was confirmed during data fitting.

### NMR spectroscopy

Caspofungin was prepared at 4 mM in the absence and presence of 1 equivalent of GaNO_3_ in 5 mm NMR tubes, total volume 600 µL 10% D_2_O in water. NMR spectra were acquired at 25 °C on a Bruker Avance III 800 spectrometer (Bruker, Rheinstetten, Germany) working at a proton operating frequency of 800.33 MHz, equipped with a 5 mm, three channel, inverse detection cryoprobe TCI-z H&F/C/N with pulse-field gradients. In 1D ^1^H spectra, solvent suppression was achieved using the excitation sculpting scheme [43]. ^1^H-^13^C correlation spectra (HSQC and HMBC) were acquired using standard Bruker pulse programs using 145 Hz and 8 Hz for the evolution of ^1^J_CH_ and ^n^J_CH_, respectively. All spectra were acquired, visualized and analyzed using TopSpin software V 3.6.

### Molecular dynamics simulations

All molecular dynamics simulations and principal component analysis (PCA) of the apo and iron-bound forms of caspofungin were performed following the overall protocol described in [44], using the GROMACS 2020.3 package [45, 46]. The initial structures of the apo and iron-bound forms of caspofungin were built using Avogadro, respecting the NMR inferred coordination geometry information for iron [47, 48] and energy minimized in GAUSSIAN09 [49]. Given that the lipophilic tail of caspofungin is not expected to play a relevant role in iron binding, we simulated caspofungin without this tail. For the iron-bound form of caspofungin, the MCPB.py tool [50] was used to streamline the parametrization procedure and define the topology of the iron coordination center, using as a starting point the structure minimized with GAUSSIAN09 [49]. The atomic partial charges for caspofungin bound to iron were calculated by a Restrained ElectroStatic Potential (RESP) fitting on electrostatic potentials calculated with GAUSSIAN09 using B3LYP and the 6-31G(d) basis set. A two-stage RESP fitting was performed with MCPB.py. The General Amber Force Field (GAFF) [51] parameters were applied to the Lennard-Jones and bonded terms. A similar procedure was used for the apo form of caspofungin, but instead of MCPB.py tool which is specific for molecules containing metals, the Antechamber framework was used to streamline the parametrization. The parameters for ions were taken from the AMBER force field [52] and the TIP3P model was used for water [53]. Each structure was inserted in a truncated dodecahedron box filled with water molecules (considering a minimum distance of 1.2 nm between protein and box walls). The total charge of the system was neutralized with the required number of Na^+^ ions, with additional Na^+^ and Cl^−^ ions added to the solution to reach an ionic strength of 0.1 M. The system was energy-minimized using the steepest descent method for a maximum of 50,000 steps with position restraints on all heavy-atoms by restraining them to the initial coordinates using a force constant of 1,000 kJ/mol in the X, Y and Z positions. Before performing the production runs, an initialization process was carried out in 5 stages of 100 ps each, and position restraints were applied to all the heavy atoms (stages 1 to 4) or to the caspofungin core (carbon and nitrogen atoms of the main chain within the cyclic peptide region) in the last stage. In the first stage, the Berendsen thermostat [54] was used to initialize and maintain the simulation at 300 K, using a temperature coupling constant of 0.01 ps, without pressure control. The second stage continued to use the Berendsen thermostat but now with a coupling constant of 0.1 ps. The third stage kept the same temperature control, but introduced isotropic pressure coupling with the Berendsen barostat [54], with a coupling constant of 5.0 ps. The fourth stage changed the thermostat to V-rescale [55], with a temperature coupling constant of 0.1 ps, and the barostat to Parrinello-Rahman [56], with a pressure coupling constant of 2.0 ps. The fifth stage is equal to the fourth stage, but position restraints were only applied on caspofungin core. For production simulations, conditions were the same as for the fifth stage, but without any restraints. In all cases, 2 fs integration steps were used. Long-range electrostatic interactions were treated with the PME [57, 58] scheme, using a grid spacing of 0.12 nm, with cubic interpolation. The neighbor list was updated every twenty steps with a Verlet cut-off with a 0.8 nm radius. All bonds were constrained using the LINCS algorithm [59]. Simulations of each system were performed over 5 µs for 8 replicates of caspofungin and 10 replicates of caspofungin plus iron (the number of replicates used for each system was based on the convergence of the system properties). The first 1 µs of each simulation was considered equilibration time, with subsequent frames used for analysis.

Principal component analysis (PCA) was applied to sets of conformational coordinates obtained from MD simulations. Before PCA, each conformation was translationally and rotationally aligned to the caspofungin core. PCs were determined using the gmx anaeig program from GROMACS [45, 46], from the entire pool of simulation trajectories, considering only the coordinates of caspofungin core atoms. The dimensionality was reduced to the two most representative PCs. Caspofungin structures for each simulation frame could then be projected as points in this two-dimensional space, enabling a simplified visual representation of the conformation space explored by caspofungin in each case. The probability density function for each trajectory projection was estimated using a gaussian kernel estimator [60, 61] implemented in LandscapeTools *get_density* software as described elsewhere [60, 62]. This procedure defines a probability density function whose values are stored for the position of each data point and for the nodes of a two-dimensional uniform grid, with a mesh size of 0.1 Å. These values were used to define an energy surface, calculated as described in [60]. The energy surface landscapes were analyzed by determining the energy minima and respective basins defined as the set of all conformations whose steepest descent path along the energy surface leads to a particular minimum [60, 63, 64]. The steepest descent paths for each grid cell were computed, with each conformation inheriting the path of its corresponding grid cell. Landscape regions with E > 6 *k*_*B*_*T* were discarded, resulting in the final set of basins for each data set.

### β-1,3-D-glucan synthase activity

Membrane fraction preparation and enzymatic activity of β-1,3-D-glucan synthase were performed as described in [65], with some modifications. *C. albicans* SC3514 cells were grown in YPD medium to mid-log phase and harvested by centrifugation at 3,000 *ɡ* for 5 min. Cells were resuspended in cold extraction buffer (50 mM Tris-HCl pH 7.5 containing 1 mM EDTA, 4 mM DTT, cOmplete Protease Inhibitor Cocktail (Roche), and 1 mM PSMF) and lysed using two cycles of French press (400 psi). The crude homogenate was centrifuged at 3,000 *g* for 10 min and the supernatant was centrifuged at 100,000 *ɡ* for 1 h. The membrane pellet was resuspended in one-tenth the original volume using extraction buffer supplemented with 33% glycerol. Total protein concentration was measured using the Bradford assay. Samples were stored at −80 °C until use. For enzyme activity assay, reactions were performed in a final volume of 50 µl containing 50 mM Tris-HCl pH 7.5, 100 µM GTP, 4 mM EDTA, 0.5% Brij 35, 6.6% glycerol, 20 mM UDP-glucose, 100 µg of membrane protein, and 0.375 µg/ml caspofungin, 500 µM FeSO_4_, or both. Reactions were incubated overnight at 25 °C and stopped by adding 10 µL of 6 M NaOH. Synthesized glucans were solubilized and quantified with alanine blue as described above.

### *Galleria mellonella* survival, fungal burden and iron quantification assays

The *Galleria mellonella* caterpillar infection model was used for the *in vivo* studies, following previously described protocols [66, 67]. *G. mellonella* larvae were reared on a diet of pollen grains and bee wax and maintained at 25 °C in darkness. For the experiments, the last instar larvae weighing 200 ± 25 mg were randomly selected. Cultures of *C. albicans* clinical isolate H19 were harvested by centrifugation and suspended in 0.9% NaCl. The concentration of *C. albicans* cells used for infection was determined based on lethality results obtained after testing increasing cell concentrations (ranging from 6 × 10^8^ cells/mL to 6 × 10^6^ cells/mL; data not shown). Stock solutions of FeCl_3_ and caspofungin were prepared in water and then diluted in 0.9% NaCl. Ten larvae per condition and per experiment were sequentially injected, via the hindmost left proleg, sanitized with 70% (v/v) ethanol, with 5 μL of (i) FeCl_3_ at either 5 mM or 1.8 mM (corresponding to 278 μM and 100 μM in the hemolymph, respectively, based on hemolymph volume of approximately 90 μL), (ii) 6 × 10^5^ *C. albicans* cells, and (iii) a caspofungin solution at 40 μg/mL (equivalent to a 1 mg/kg dose, assuming an average larval weight of ∼ 200 mg). The injections were performed with a 30-minute interval between each. As control, a set of larvae was injected with the same volume of 0.9% NaCl. The larvae were placed in Petri dishes and stored in the dark at 37 °C for 3 days, with survival monitored at 24-h intervals. Larvae were considered dead if they showed no movement in response to touch. Kaplan-Meier survival curves were generated from three independent experiments. For fungal burden assessment, hemolymph was collected from live larvae after a 48 h incubation period. Ten larvae were anesthetized on ice, surface-sterilized with 70% (v/v) ethanol and punctured in the abdomen with a sterile needle. The outflowing hemolymph was immediately collected, and 10 μL were plated onto YPD plates supplemented with 100 μg/mL chloramphenicol. Plates were incubated at 30 °C for 24 h, and CFUs were counted. For iron quantification, two groups of 12 larvae were injected with either 5 μL of FeCl_3_ at 5 mM or 5 μL of 0.9% NaCl (control). After 48 h, hemolymph was collected as described above into tubes containing a few crystals of phenylthiourea to prevent melanization and stored at −20 °C. Samples were centrifuged at 20,000 *ɡ* for 5 min, and the concentration of total free iron was measured using the Iron Assay Kit (Sigma-Aldrich^®^) according to the manufacturer’s instructions.

## 3. Results

### 3.1. Iron antagonizes the activity of caspofungin against *C. albicans* and other medically relevant fungi

The sensitivity phenotype of *C. albicans* to caspofungin is attenuated when the growth medium is supplemented with iron(II) or iron(III) (Fig. 1A). Consistent with this observation, flow cytometry with propidium iodide (PI), a membrane impermeable dye typically excluded from viable cells, revealed that the combination of ferrous iron and caspofungin strongly reduced PI staining by more than 50% compared to caspofungin treatment alone (Fig. 1B).

**Figure 1.**
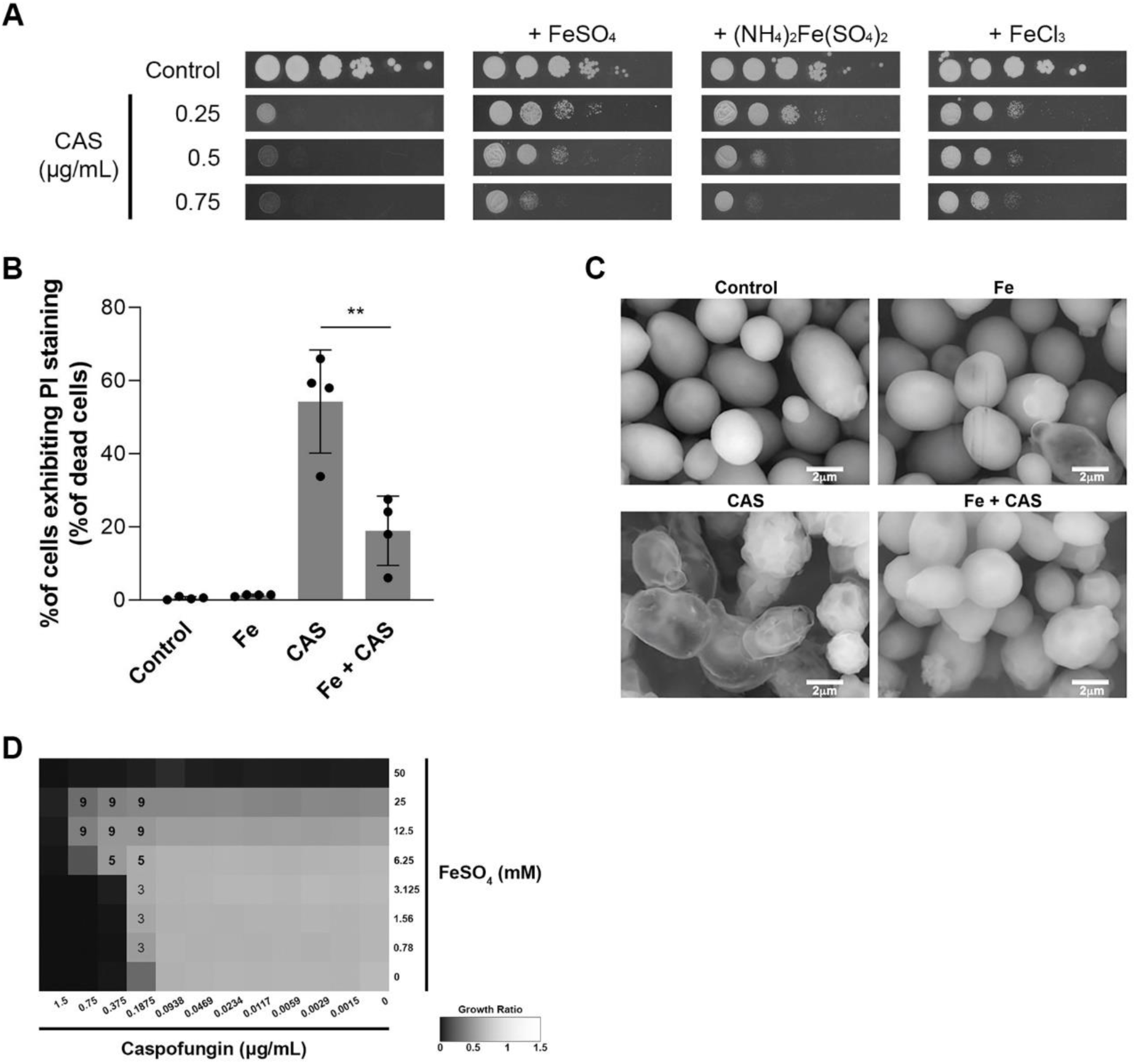
Iron supplementation alleviates caspofungin in *C. albicans*. **(A)** Growth sensitivity of *C. albicans* SC5314 was assessed on SC agar plates (Control) containing the indicated concentrations of caspofungin (CAS), different sources of iron, or their combination. Growth was recorded after 48 h of incubation at 30 °C. **(B)** Detection of dead cells using flow cytometry with propidium iodide staining (PI) under control conditions or following treatment with FeSO_4_ (Fe), caspofungin (CAS) or both (Fe + CAS) for 24 h. Statistical significance of differences was calculated using Student’s *t-*test (** P < 0.001). **(C)** SEM images of *C. albicans* cells left untreated (Control), or treated with FeSO_4_ (Fe), caspofungin (CAS) or their combination (Fe + CAS) for 24 h. **(D)** Checkerboard assays were conducted to evaluate *C. albicans* growth under combinations of FeSO_4_ (ranging from 50 to 0.78 mM) and caspofungin (ranging from 1.5 to 0.0015 µg/mL). Growth ratios were assessed after 24 h at 30 °C by measuring OD_600_ and normalizing the values relative to untreated controls (gradient bar). The fractional inhibitory concentration index (ƩFIC) was calculated for the specified combinations of caspofungin and FeSO_4_. Combinations exhibiting antagonism are highlighted with bold ƩFIC values.

Caspofungin inhibits the synthesis of β-1,3-D-glucans, which are major structural components of the fungal cell wall [68, 69], causing osmotic instability that leads to cell swelling and lysis [70]. Scanning electron microscopy (SEM) was used to assess the impact of iron and caspofungin on *C. albicans* cell morphology. *C. albicans* cells have a well-defined oval shape with a smooth surface (Fig. 1C). Treatment with iron did not affect cell morphology. In contrast, caspofungin-treated cells exhibited a rough and irregular surface, with most cells appearing translucent, indicating they were empty due to cell wall disruption, lysis, and the release of intracellular contents. However, cells treated with both iron and caspofungin, although still displaying some surface irregularities, appeared intact, suggesting that iron prevents cell wall disruption caused by caspofungin.

The interaction between caspofungin and iron was further examined using checkerboard microdilution assays. The effect of all possible combinations of two-fold serial dilutions of caspofungin (1.5 to 0.0015 µg/mL) and iron (50 to 0.78 mM) on *C. albicans* growth was evaluated (Fig. 1D). The MIC_Fe_ was 50 mM and the MIC_CAS_, was 0.1875 µg/mL. Antagonism between iron and caspofungin (FIC_Fe_ + FIC_CAS_ > 4) was observed for iron concentrations equal or greater than 6.25 mM. The MIC_CAS_ increased 4-fold and 8-fold when iron was added at a concentration of 6.25 mM and 12.5 mM, respectively. Although iron does not exhibit a true antagonism with caspofungin at concentrations below 6.25 mM, an increase in the MIC_CAS_ was still observed (FIC_Fe_ + FIC_CAS_ = 3) at the lowest iron concentration tested (0.78 mM) (Fig. 1D). Remarkably, iron supplementation at concentrations as low as 31.25 µM reduced the toxicity of caspofungin (Fig. S1). These findings indicate that even low iron levels confer a growth advantage to *C. albicans* cells challenged with caspofungin.

The antagonism between iron and caspofungin was also evident in *C. albicans* clinical isolates (Table S2). The MIC_CAS_ across all isolates ranged from 0.0469 to 0.375 µg/mL, and the MIC_Fe_ was 50 mM except for 5 isolates, for which MIC_Fe_ could not be determined (> 50 mM). Among the 19 clinical isolates tested, 12 demonstrated clear antagonism between iron and caspofungin (Table S2).

To assess whether other transition metals (cobalt, nickel, copper, and zinc) affect caspofungin activity against *C. albicans*, similar experiments were performed, but no antagonistic effect (FIC_Metal_ + FIC_CAS_ > 4) was observed (Fig. S2).

Checkerboard dilution assays on *Saccharomyces cerevisiae* and six clinically relevant fungal species (*C. glabrata*, *C. parapsilosis*, *C. tropicalis*, *C. dubliniensis*, *C. krusei*, and *Aspergillus fumigatus*) revealed that the antagonism between iron and caspofungin also occurred in *C. dubliniensis* and *C. tropicalis* (Fig. S3), the yeast species most closely related to *C. albicans* [71], and in *A. fumigatus*, where the MEC_CAS_ increased 8-fold with 6.25 mM FeSO_4_ (Fig. S4).

Further testing with other antifungal drugs (micafungin, anidulafungin, fluconazole, and amphotericin B) showed no true antagonism with iron, indicating that the effect is specific to caspofungin (Fig. S5).

### 3.2. Iron overload reduces the ability of caspofungin to prevent *C. albicans* biofilm formation

In *C. albicans*, biofilm formation is a major virulence factor that contributes to the severity of infections [6]. Biofilm development can be divided into four stages: adherence, initiation (or proliferation), maturation and dispersion [6]. The potential impact of iron on the antibiofilm activity of caspofungin was investigated at different stages of biofilm formation. Thus, three biofilm viability assays were performed, where both iron and caspofungin were (i) present throughout the assay (biofilm sustained inhibition assay), (ii) added after the adhesion phase (biofilm developmental inhibition assay), and (iii) added after the formation of a mature biofilm (mature biofilm disruption assay), as shown in Fig. 2A.

**Figure 2.**
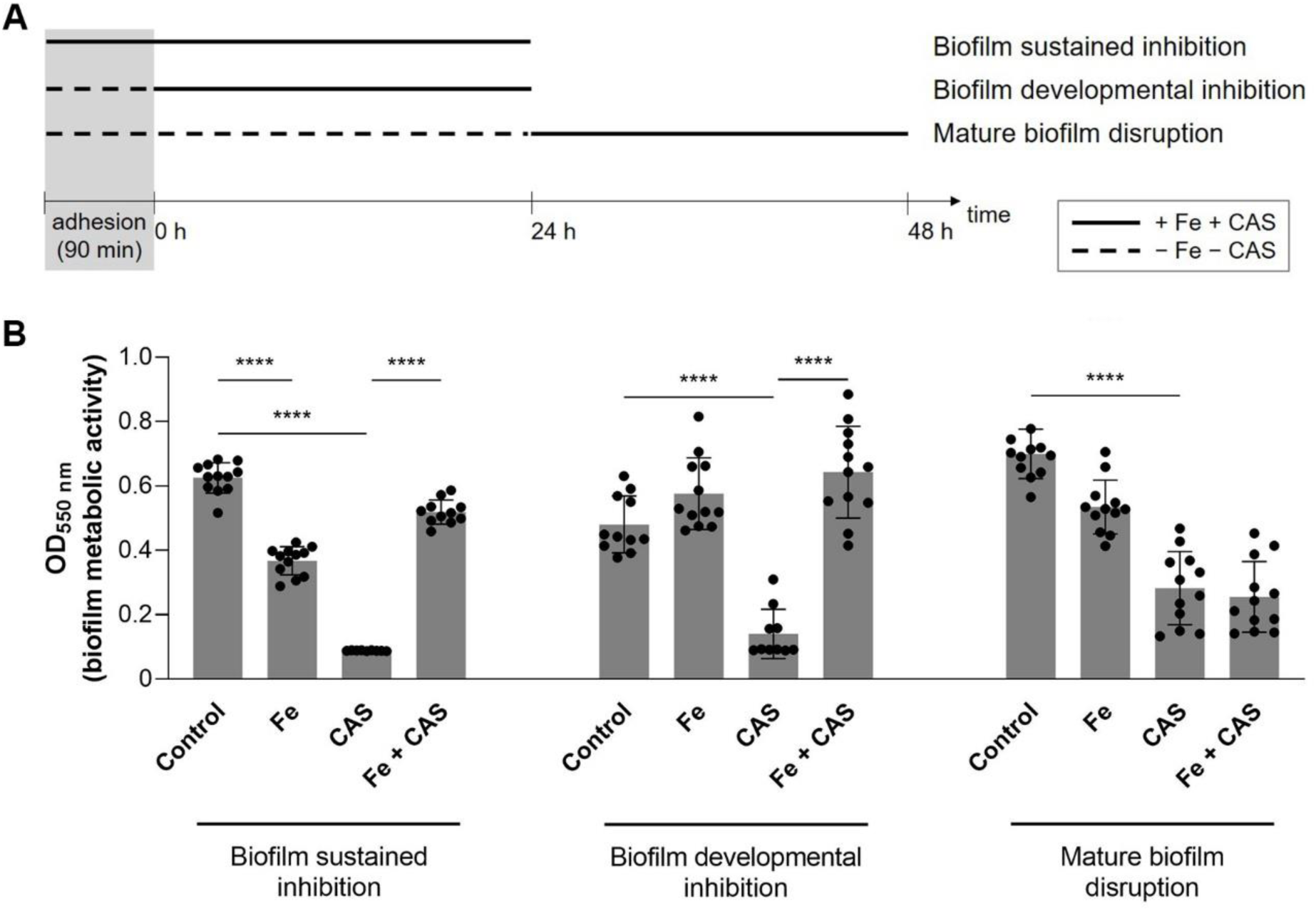
Inhibition of biofilm formation by caspofungin is prevented by iron. **(A)** Schematic representation of the procedures used for the biofilm assays performed in this study. **(B)** Biofilm viability of *C. albicans* was assessed for untreated biofilms (Control) or biofilms treated with 0.5 mM FeSO_4_ (Fe), caspofungin (CAS) at concentrations of 0.0469 µg/mL, 0.0938 µg/mL, 0.0469, µg/mL or 0.1875 µg/mL (corresponding to assays for sustained inhibition, developmental inhibition and mature biofilm disruption, respectively), or a combination of both compounds (Fe + CAS). Biofilm viability was quantified using the MTT colorimetric assay. Statistical significance of differences was determined using Student’s *t-*test (**** P < 0.0001; * P < 0.05).

The sessile minimum inhibitory concentrations of caspofungin (SMIC_CAS_) were first determined as follows: 0.0469 µg/mL for sustained inhibition, 0.0938 µg/mL for developmental inhibition, and 0.1875 µg/mL for mature biofilm disruption. These caspofungin concentrations were combined with iron in the corresponding biofilm viability assays (Fig. 2A). Biofilm viability was assessed using the 3-(4,5-dimethyl-2-thiazolyl)-2,5-diphenyl-2H-tetrazolium bromide (MTT) assay, in which metabolically active cells reduce MTT to a dark purple formazan product [37].

Interestingly, iron alone decreased biofilm viability in the sustained inhibition assay. In all assays, biofilm viability was significantly higher in the presence of iron combined with caspofungin (Fe + CAS) compared to caspofungin alone, except in the mature biofilm disruption assay, where neither ferrous (Fig. 2B) nor ferric (Fig. S6) iron had an effect. These results suggest that iron antagonizes the activity of caspofungin during the early stages of biofilm formation, but not after biofilm maturation.

### 3.3. The effect of iron on caspofungin activity is independent of changes in the composition of *C. albicans* cell wall

Previous studies have shown that in *C. albicans*, iron modulates the levels of cell wall components, making the yeast resistant to cell wall-perturbing agents [31, 72]. Therefore, we first hypothesized that the observed antagonism between caspofungin and iron (Fig. 1) could be a result of changes in cell wall composition induced by iron. To explore this hypothesis, the effect of iron on *C. albicans* major cell wall components (chitin, β-1,3-D-glucans, and mannans) was analyzed using confocal microscopy and flow cytometry. Under our experimental conditions, iron alone did not affect cell wall composition (Fig. 3A and B). Supporting this observation, overnight treatment with iron prior to exposure to cell wall-perturbing agents (caspofungin, calcofluor white and tunicamycin) did not confer resistance to these stressors (Fig. 4A). As expected, β-1,3-D-glucan levels decreased following caspofungin treatment, while chitin levels greatly increased (Fig. 3B), reflecting a compensatory mechanism to counteract the reduced β-1,3-D-glucans in the cell wall [73]. Iron appeared to mitigate these alterations, restoring the levels of β-1,3-D-glucans and chitin to nearly normal levels (Fig. 3B). Interestingly, the combination of iron and caspofungin resulted in increased mannan levels (Fig. 3B); however, this is unlikely to explain the observed antagonistic effect between the metal and the drug. This conclusion is supported by the observation that the *C. albicans* mutant strain ΔΔ*mnn10*, which exhibits truncated *N*-linked oligosaccharides and glycosylation defects in mannans [74], displayed the same MIC_CAS_ and degree of antagonism between caspofungin and iron as its parental counterpart (Fig. S7).

**Figure 3.**
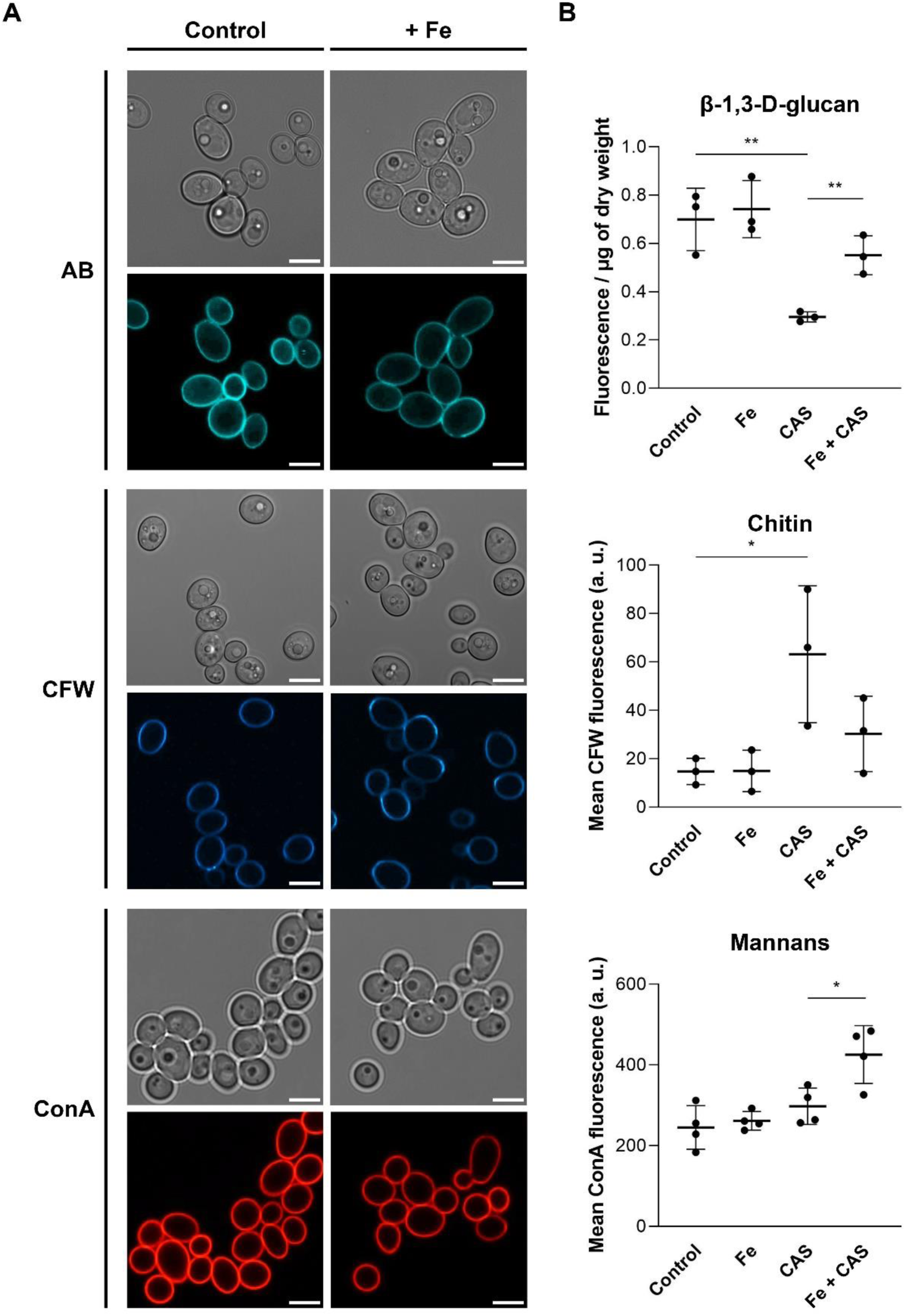
Alterations in the cell wall composition of *C. albicans* do not account for the antagonistic effect of iron on caspofungin activity. **(A)** Confocal microscopy images of *C. albicans* SC5314 cells left untreated (Control) or treated with FeSO_4_ (+ Fe), and stained with calcofluor white (CFW), aniline blue (AB) or concanavalin A (ConA), to visualize chitin, β-1,3-D-glucans, and mannans, respectively. Scale bar: 5 μm. **(B)** Quantification of cell wall components of *C. albicans* left untreated (Control), or treated overnight with FeSO_4_ (Fe), caspofungin (CAS) or a combination of both (Fe + CAS). Statistical significance was determined using Student’s *t*-test (** P < 0.01; * P < 0.05).

**Figure 4.**
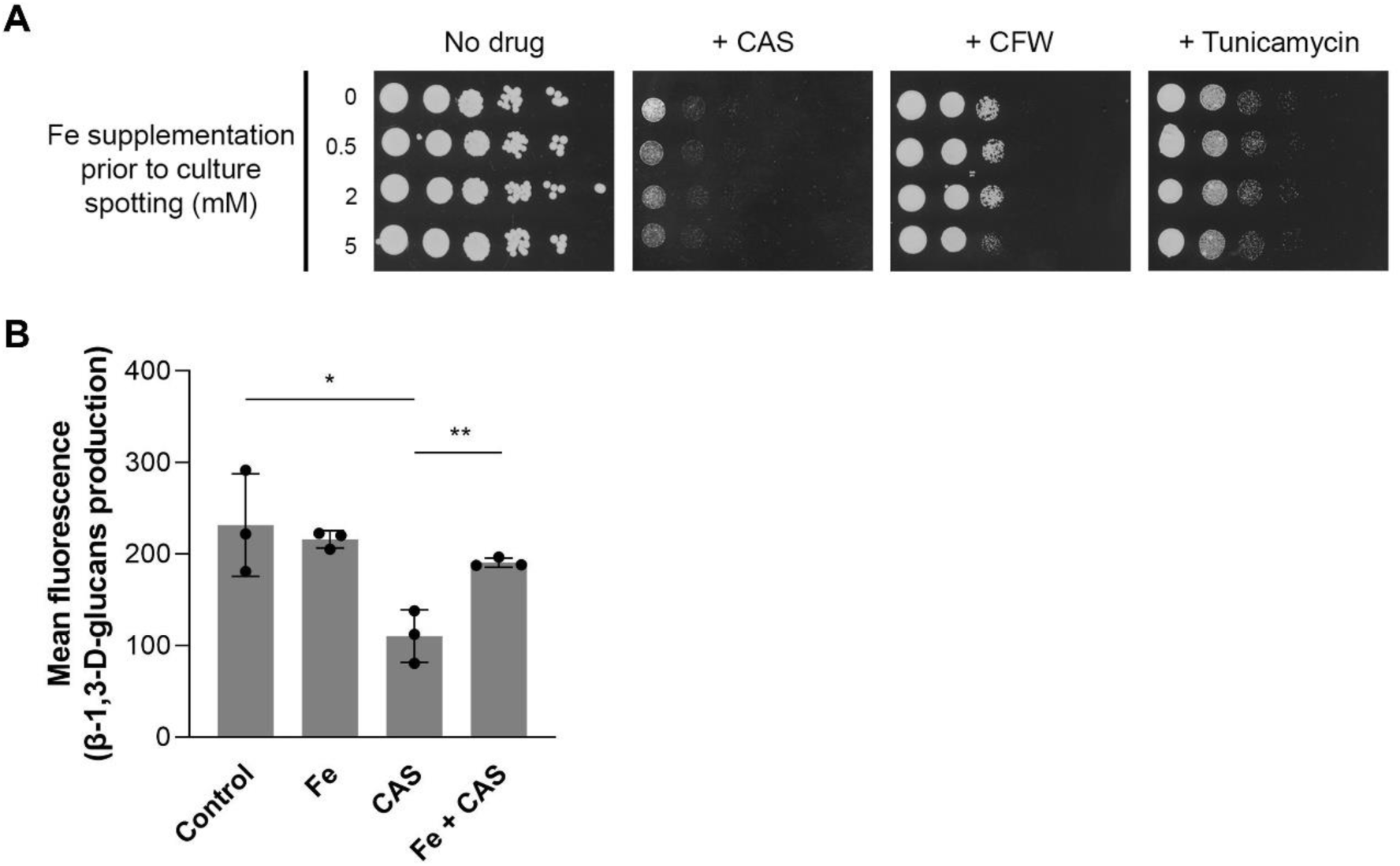
The antagonistic effect between iron and does not appear to be dependent on cellular metabolism. **(A)** Growth sensitivity of *C. albicans* SC5314 cells on SC agar plates containing 0.5 µg/mL caspofungin (+ CAS), 50 µg/mL calcofluor white (+ CFW) and 1.5 µg/mL tunicamycin. Cells were grown in the indicated concentrations of FeSO_4_ before being spotted onto the agar plates. Growth was recorded after 48 h at 30 °C, except for caspofungin which was recorded after 72 h. **(B)** *C. albicans* membrane fractions (containing β-1,3-D-glucan synthase) were left untreated (Control) or treated with FeSO_4_ (Fe), caspofungin (CAS) or both (Fe + CAS). Significance of differences was calculated using the Student’s *t*-test (** P < 0.01; * P < 0.05).

To investigate whether iron directly interferes with caspofungin, the *in vitro* activity of β-1,3-D-glucan synthase was assessed in the presence of caspofungin, iron, or both (Fig. 4B). β-1,3-D-glucan synthase activity was reduced by caspofungin but restored in the presence of both caspofungin and iron, consistent with the observation that cell wall β-1,3-D-glucan levels were also restored *in vivo* when iron and caspofungin were combined (Fig. 3B).

### 3.4. Iron binding induces conformational changes in caspofungin

We next sought to understand the molecular basis underlying the reduced inhibition of β-1,3-D-glucan synthase by caspofungin in the presence of iron. Given that caspofungin contains several chemical groups capable of coordinating metals, such as amides, amines and hydroxyls, we hypothesized that caspofungin may bind to iron. To investigate this, an indirect method using UV-Vis spectroscopy with thiocyanate as an iron(III) indicator was employed (Fig. 5A). When bound to thiocyanate, iron(III) displays a maximum absorbance centered around 450 nm. The addition of 1.2 equivalents of caspofungin led to an accentuated decrease in this absorbance, indicating that caspofungin has a higher affinity for iron(III) than the indicator and is able to displace the metal from the thiocyanate-iron(III) complex. Although a decrease in the absorbance was also observed with the addition of 1.2 equivalents of micafungin, it was less accentuated than for caspofungin (Fig. 5B). To confirm this qualitative assessment, the conditional binding and dissociation constants of both echinocandins to iron(III) were determined (Fig. S8). Indeed, iron(III) binding was found to be *ca.* two orders of magnitude higher for caspofungin compared to micafungin (log *K*_f_ of 5.71 *vs.* 3.93, Table 1).

**Figure 5.**
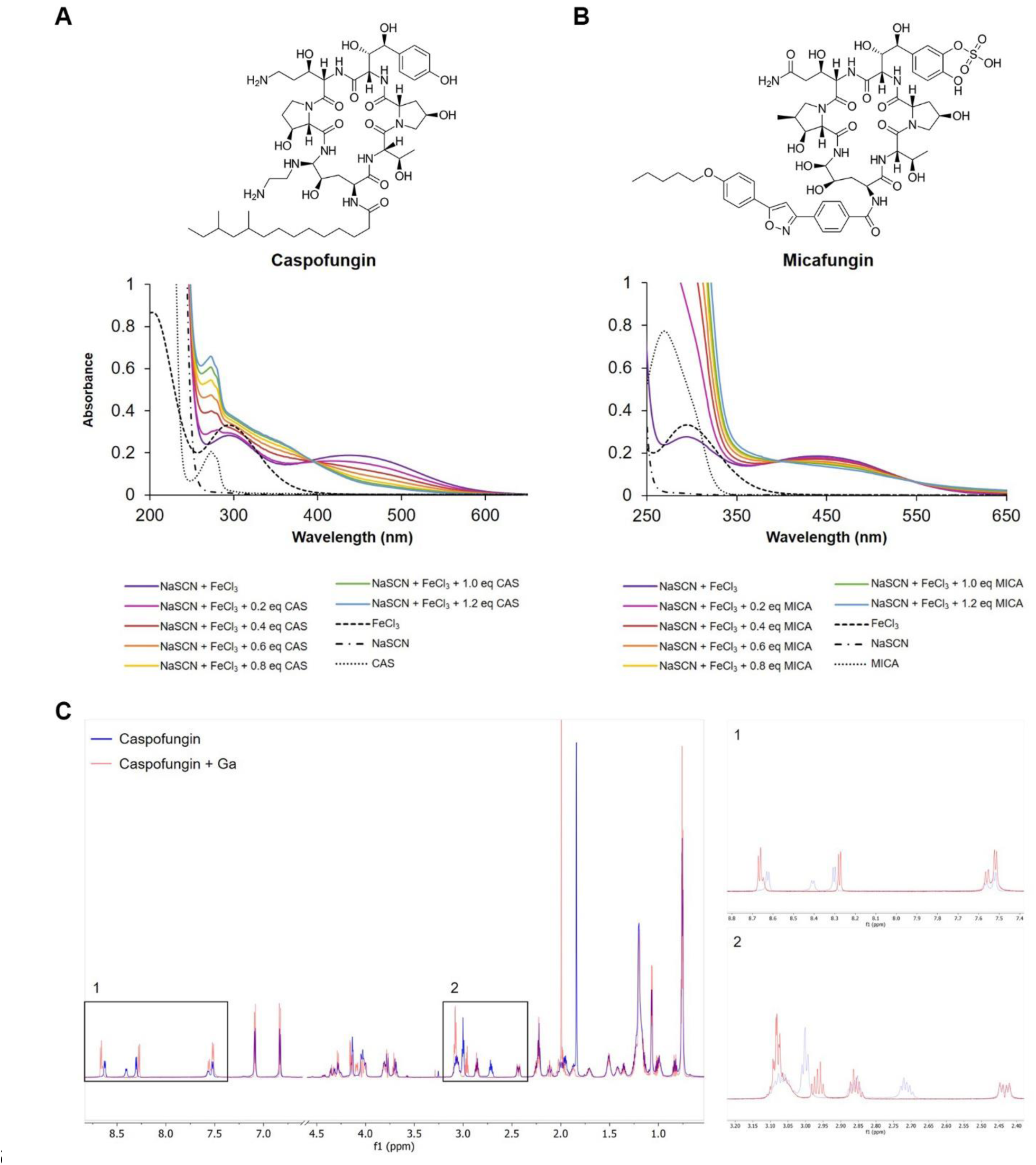
Caspofungin binds iron. UV-Vis absorption spectra depicting the competition for iron(III) between thiocyanate (NaSCN) and **(A)** caspofungin (CAS) or **(B)** micafungin (MICA). Absorption profiles were recorded after stepwise addition of the echinocandins up to 1.2 equivalent into a solution containing 7.5 mM NaSCN and 150 µM FeCl_3_. **(C)** ^1^H-NMR spectra of caspofungin (4 mM) in the absence (blue) or the presence (red) of 4 mM Ga(NO_3_)_3_. The central section of the spectrum between 4.6 and 6.5 ppm contained no signals apart from the residual water line and were removed.

**Table 1.**
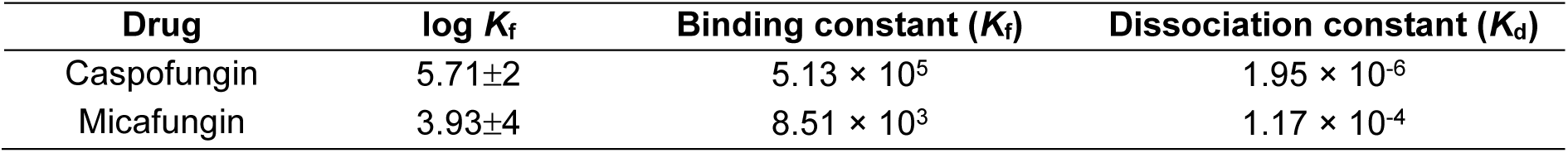
Conditional iron binding and dissociation constants for caspofungin and micafungin.

To gain further insight into iron binding to caspofungin, we compared the shifts in the NMR signals of caspofungin in the absence and presence of 1 equivalent of gallium(III), a diamagnetic iron(III) surrogate with similar coordination properties. The assignment of NMR signals (both in ^1^H and ^13^C) was achieved by comparison of the HSQC and HMBC maps with the values reported in the literature [75]. While the majority of the signals in the NMR spectra remained unaltered after the addition of gallium(III), a few distinct signals displayed noticeable shifts, particularly those attributed to the ethylenediamine moiety (δ^1^H=2.3–3.2) and two of the amide groups (δ^1^H=7.4–8.7) of caspofungin (Fig. 5C). These findings strongly suggest that caspofungin forms a complex with gallium(III), coordinated by the two amine and two amide groups, and it is expected that a similar complex would form with iron(III).

To test whether iron binding could affect the caspofungin conformation, and eventually impact its binding to β-1,3-D-glucan synthase, molecular dynamics (MD) simulations were performed to investigate the structural dynamics of caspofungin when bound to iron as compared to the unbound drug. PCA was applied to the frames extracted from the trajectory obtained from MD simulations, which revealed two conformations corresponding to the lowest energy structures for both systems (Fig. 6). For unbound caspofungin, we observed two deep basins (basin 0 and basin 1). A closer analysis of caspofungin conformations corresponding to each basin revealed that these conformations are similar to one another (which is consistent with the fact that they are located in proximal regions of the energy landscape), with some differences mainly in the side chains. The energy barrier between these conformations is quite shallow and one can think of this system as having a single conformational state that contains two similar sub-states (corresponding to the two basins). In contrast, the basins of caspofungin bound to iron correspond to conformations that are quite distinct from those of unbound caspofungin (Fig. 6). The two conformations observed in caspofungin:iron system are also quite distinct from one another, which is reflected in the fact that these basins are located in distinct regions of the energy landscape. In spite of their differences, both conformations showed that the caspofungin core is closed over itself when bound to iron, with this effect being more pronounced in basin 1. Overall, these results demonstrate that iron binding significantly alters the conformation of caspofungin.

**Figure 6.**
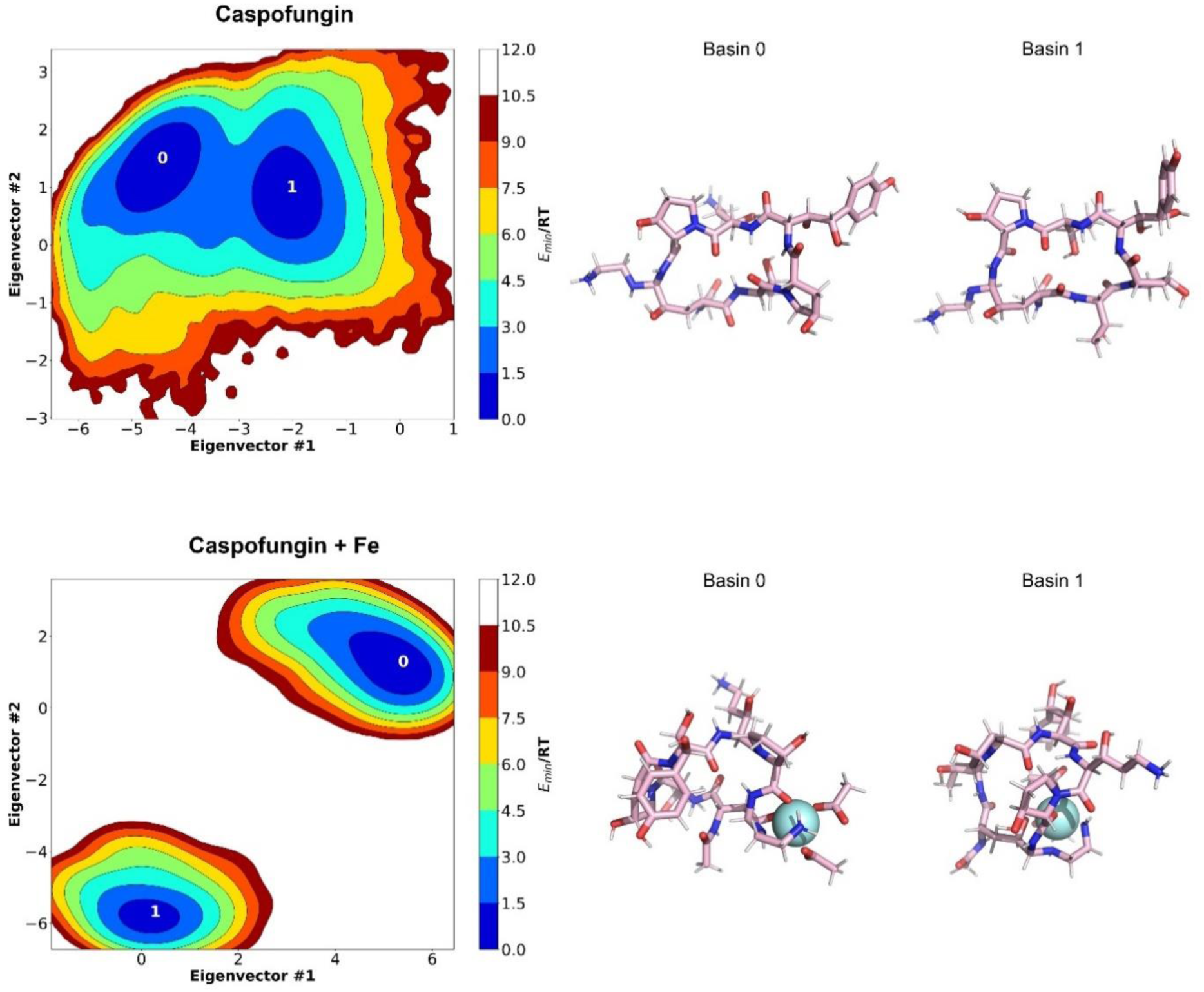
Caspofungin undergoes conformational changes in the presence of iron. Plots of two-dimensional PCA of caspofungin conformational dynamics in water, comparing caspofungin in the unbound state and bound to iron. Color scale represents k_B_T, and basins with k_B_T < 3 are numbered. Snapshots of the lowest energy conformations are shown for each system.

### 3.5. Iron overload reduces the *in vivo* activity of caspofungin against *C. albicans* in the *Galleria mellonella* infection model

*In vivo* experiments were performed to assess the effect of iron overload on caspofungin treatment in *Galleria mellonella* larvae infected with a *C. albicans* clinical isolate. Four groups of larvae were used (Fig. 7): a group infected with *C. albicans* (group *C. albicans*), a group preloaded with iron and infected with *C. albicans* (*C. albicans* + Fe), a group infected with *C. albicans* and treated with caspofungin (*C. albicans* + CAS), and a group preloaded with iron, infected with *C. albicans* and treated with caspofungin (*C. albicans* + Fe + CAS). Neither iron nor caspofungin was toxic to the larvae at the concentrations used in these *in vivo* assays, specifically 280 μM of FeCl_3_ or 1 mg/kg of caspofungin in the hemolymph of larvae (data not shown). None of the larvae in either the *C. albicans* or *C. albicans* + Fe groups survived at 72 h. When treated with caspofungin, the survival rate of infected larvae in the *C. albicans* group was 60%. However, this survival rate dropped significantly to 16.7% when larvae were preloaded with iron (*C. albicans* + Fe + CAS, Fig. 7A). The same effect was observed upon larvae preloading with lower concentrations of iron (100 μM FeCl_3_ in the hemolymph of larvae, Fig. S9). As expected, the fungal burden in larvae injected with both iron and caspofungin (*C. albicans* + Fe + CAS) was higher than in those injected with caspofungin alone (*C. albicans* CAS) (Fig. 7B). Moreover, we confirmed that iron levels in the lymph of larvae injected with iron remained increased for at least up to 48 h (Fig. 7C). In summary, these results suggest that iron overload may adversely affect the treatment of *C. albicans* infections with caspofungin.

**Figure 7.**
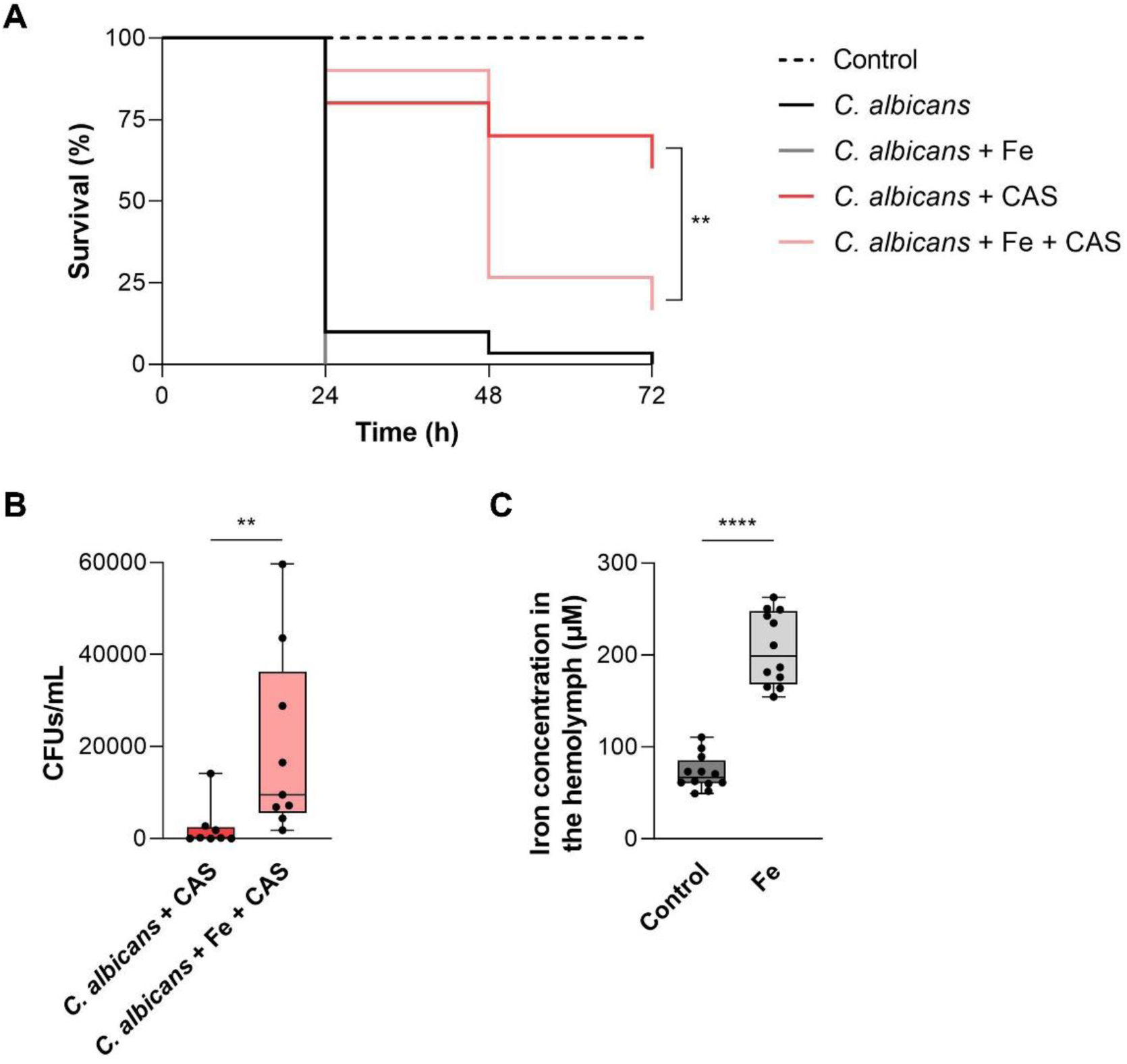
Iron overload impairs the *in vivo* activity of caspofungin against *C. albicans* in a *G. mellonella* infection model. **(A)** Survival rates of *G. mellonella* larvae injected with 0.9% NaCl (Control), infected with the *C. albicans* clinical strain H19 (*C. albicans*), preloaded with 280 μM FeCl_3_ and infected with H19 (*C. albicans* + Fe), infected with H19 and treated with 1 mg/kg caspofungin (*C. albicans* + CAS), or preloaded with 280 μM FeCl_3_, infected with H19 and treated with 1 mg/kg caspofungin (*C. albicans* + Fe + CAS). Statistical significance of differences in survival rates was determined using the log-rank test (*** P < 0.001; ** P < 0.01). **(B)** Fungal burden in the hemolymph of infected *G. mellonella* treated with 1 mg/kg caspofungin. Prior to infection, larvae were treated with 280 μM FeCl₃ (*C. albicans* + Fe + CAS) or left untreated (*C. albicans* + CAS). Hemolymph was collected from infected larvae after 48 h and plated on YPD agar plates for CFU counting. **(C)** Hemolymph iron levels of *G. mellonella* larvae injected with 0.9% NaCl (Control) or with 280 μM FeCl_3_ (Fe) after 48 h. Statistical significance of differences was calculated using Student’s *t*-test (**** P < 0.0001; ** P < 0.01; * P < 0.05).

## 4. Discussion

IFIs have been on the rise, posing a significant health threat, particularly to hospitalized and immunocompromised patients [3, 4]. Iron overload medical conditions have been shown to exacerbate these infections [27–29] and are associated with poor clinical outcomes [23]. Recently, iron overload has been found to affect the *in vitro* susceptibility of *C. albicans* to several antifungals [26], suggesting that such condition might also impact the treatment of infections caused by this fungal species.

In this study, we show that iron overload impairs not only the *in vitro*, but also the *in vivo* antifungal activity of caspofungin (Fig. 1 and 7), the preferred treatment for invasive candidiasis [10]. Our results provide evidence of a true antagonistic effect between the metal and the drug (Fig. 1), which is also observed in *C. albicans* clinical isolates (Table S2), and other clinically relevant fungal species, including *C. tropicalis*, *C. dubliniensis* and *A. fumigatus* (Fig. S3 and S4), broadening the potential impact our observation. The intriguing finding that the antagonistic effect is not observed across all *Candida* species led us to hypothesize that interspecies variations in the structural features of the β-1,3-D-glucan synthase catalytic subunit (Fks) may account for this observation. In this context, since the exact binding mechanism of caspofungin to Fks remains unknown, the MD studies conducted here provide valuable clues by suggesting that iron binding may promote a change in the conformational equilibrium of caspofungin, which, in turn, may disturb its interaction with Fks.

Importantly, the antagonistic effect between iron and caspofungin was also observed in *C. albicans* biofilms, a well-known critical virulence factor [76] (Fig. 2). Interestingly, caspofungin inhibition of *C. albicans* biofilms was reversed by iron, at different stages of biofilm formation, except in mature biofilms (Fig. 2B and S6). Mature biofilms consist of cells surrounded by an extracellular matrix composed of proteins, carbohydrates, lipids and nucleic acids, which serves as a protective physical barrier against environmental factors and provides structural integrity to the biofilm [6]. Thus, one possible explanation for this dissonant phenotype is that the extracellular matrix may contain molecules that adsorb or chelate iron, thereby impeding its antagonistic effect with caspofungin. In fact, it has been reported that mature biofilms of the Gram-positive bacterium *Bacillus subtilis* secrete pulcherriminic acid, an iron chelator that halts biofilm expansion by depleting iron in the surrounding environment. This mechanism enables *B. subtilis* to defend its niche against neighboring bacteria [77]. The yeast *Metschnikowia pulcherrima* also produces pulcherrimin, which is associated with defense against other species [78]. Although it is not known whether *C. albicans* biofilms produce similar compounds, the presence of other molecules in the extracellular matrix capable of binding iron and neutralizing its antagonism with caspofungin cannot be ruled out. An alternative hypothesis could be that if the effect were iron(III)-dependent, the hypoxic environment of mature biofilms [79] might hinder the oxidation of iron(II), which would not antagonize caspofungin. However, this is unlikely, as a similar phenotype was observed with ferric iron (Fig. S6). Lastly, the finding that the addition of iron from the initial stages of biofilm formation (adhesion stage) decreased biofilm viability (Fig. 2) may be explained by the fact that under iron overload conditions *C. albicans* genes encoding proteins involved in cell wall remodelling and adhesion, both essential for biofilm formation, are overall repressed [80].

Recent studies have suggested that iron supplementation alters the *C. albicans* cell wall composition, leading to increased resistance to cell wall perturbing agents, including caspofungin [26, 31]. Contrary to these reports, we observed that the antagonistic effect between iron and caspofungin persists even under conditions where cell wall composition remains unchanged (Fig. 3). Furthermore, this effect is independent of the impact of high iron on cellular metabolism, as pre-treatment with iron did not protect yeast cells from caspofungin and other cell wall perturbing agents (Fig. 4A). Consistent with these findings, and suggesting a direct effect of iron on caspofungin, iron prevents the *in vitro* inhibition of β-1,3-D-glucan synthase when combined with caspofungin, but not when used alone (Fig. 4B). The discrepancy with previous studies may arise from differences in experimental conditions. In those studies, cells were adapted to low iron (absence of iron in the medium) and high iron (100 µM iron added to iron-depleted medium) for extended periods [26], whereas we supplemented iron-replete medium with higher iron concentrations for shorter periods. It is possible that under the former conditions, both mechanisms coexist. Indeed, this could explain why in those studies some clinical *C. albicans* strains, which showed no cell wall composition alterations, continued to exhibit the antagonistic effect between iron and caspofungin [26].

UV-Vis and NMR studies on caspofungin in the presence and absence of ferric iron revealed that caspofungin binds iron (Fig. 5). The ethylenediamine moiety and two of the amide groups of caspofungin were identified as being involved in the binding of gallium(III), a diamagnetic iron(III) surrogate with similar coordination properties (Fig. 5C), suggesting that a similar coordination is likely observed with iron. The ethylenediamine moiety is absent in the other echinocandin drugs, which may explain the lack of antagonism between iron and micafungin (Fig. S5), as well as why micafungin does not bind iron as effectively as caspofungin (Fig. 5A and B). These findings, together with the impact of iron on the inhibitory activity of caspofungin on β-1,3-D-glucan synthase (Fig. 4B) and the pronounced conformational change of caspofungin upon iron binding predicted by MD simulations (Fig. 6), strongly support the hypothesis that the caspofungin-iron complex is unable to efficiently inhibit *C. albicans* β-1,3-D-glucan synthase.

In the bloodstream, iron bioavailability is extremely low due to its sequestration by transferrin, a protein with high affinity for iron [81]. However in patients with heredity hemochromatosis, the concentration of chelatable iron, or non-transferrin bound iron (NTBI), in the plasma can reach up to 16.3 µM [82]. Our *in vitro* experiments showed that a true antagonism between iron and caspofungin in *C. albicans* is only observed at iron concentrations far exceeding physiological levels (Fig. 1D). However, lower iron concentrations were still able to reduce the *in vitro* efficacy of caspofungin against *C. albicans* (Fig. 1D and S1). Supporting the impact of lower iron concentrations on caspofungin activity, *in vivo* experiments using *G. mellonella* clearly demonstrate that loading larvae with iron at a concentration of 100 μM in the hemolymph is sufficient to compromise caspofungin activity against *C. albican*s (Fig. S9). Given that insect hemolymph contains transferrin, which is typically only 30–40 % saturated with iron [83, 84], the concentration of NTBI in larvae hemolymph is anticipated to be well below 100 μM, supporting the idea that the antagonistic effect between caspofungin and iron may also occur at biologically achievable iron concentrations.

Our results underscore that under medical conditions of iron overload, caspofungin may lose efficacy against invasive infections caused by *C. albicans*. Since the antagonism between iron and caspofungin is also evident, albeit to a lesser extent, in other clinically relevant fungi, our findings may potentially be extended to fungal infections caused by other species.

## 5. Acknowledgements

We are grateful to Prof. Neta Dean (Stony Brook University) for kindly providing the *C. albicans* BWP17 and SKY68 strains, and Prof. Paulo Paixão (NOVA Medical School, Universidade Nova de Lisboa; SYNLAB, Hospital da Luz) for supplying the *C. albicans* clinical isolates. We also thank Dr. Carolina Feliciano for her assistance with confocal microscopy.

