## Supplemental files for "Caspofungin binding to iron compromises its antifungal efficacy against *Candida albicans*"

**Table S1. Fungal strains used in this study.**

| Strain | Genotype / Description | Source |
|---|---|---|
| *Candida albicans* SC5314 | Wild-type strain | ATCC |
| *Candida albicans* BWP17 | ura3Δ::limm434/ura3Δ::limm434 his1::hisG/his1::hisG arg4::hisG/arg4::hisG | [1] |
| *Candida albicans* SKY68 | BWP17 mnn10Δ::ARG4/mnn10Δ::HIS1 | [1] |
| *Candida dubliniensis* MYA-577 | Wild-type strain | ATCC |
| *Candida tropicalis* 7501 | Wild-type strain | ATCC |
| *Candida parapsilosis* 22019 | Wild-type strain | ATCC |
| *Candida glabrata* 2001 | Wild-type strain | ATCC |
| *Saccharomyces cerevisiae* BY4742 | Wild-type strain | Euroscarf |
| *Candida krusei* 6258 | Wild-type strain | ATCC |
| *Aspergillus fumigatus* (AF1160+) | akuB (KU80)-delta pyrG1 MAT1-1 | – |
| H2 | *Candida albicans* clinical strain | vaginal mucosa |
| H3 | *Candida albicans* clinical strain | vaginal mucosa |
| H4 | *Candida albicans* clinical strain | vaginal mucosa |
| H5 | *Candida albicans* clinical strain | vaginal mucosa |
| H7 | *Candida albicans* clinical strain | vaginal mucosa |
| H8 | *Candida albicans* clinical strain | vaginal mucosa |
| H9 | *Candida albicans* clinical strain | vaginal mucosa |
| H10 | *Candida albicans* clinical strain | vaginal mucosa |
| H11 | *Candida albicans* clinical strain | vaginal mucosa |
| H12 | *Candida albicans* clinical strain | vaginal mucosa |
| H13 | *Candida albicans* clinical strain | vaginal mucosa |
| H14 | *Candida albicans* clinical strain | ascitic liquid |
| H16 | *Candida albicans* clinical strain | vaginal mucosa |
| H18 | *Candida albicans* clinical strain | vaginal mucosa |
| H19 | *Candida albicans* clinical strain | vaginal mucosa |
| H21 | *Candida albicans* clinical strain | vaginal mucosa |
| H22 | *Candida albicans* clinical strain | vaginal mucosa |
| H23 | *Candida albicans* clinical strain | ascitic liquid |
| H24 | *Candida albicans* clinical strain | urine |

**Table S2. Iron acts antagonistically with caspofungin in *C. albicans* clinical isolates.**

| Clinical isolate | $MIC_{CAS}$ (µg/mL) | $MIC_{Fe}$ (mM) | Highest ΣFIC |
|---|---|---|---|
| H2 | 0.375* | 50* | 5* |
| H3 | 0.0938 | 50 | 5 |
| H4 | 0.0938 | 50 | 3 |
| H5 | 0.0938 | 50 | 5 |
| H7 | 0.0938 | 50 | 2.5 |
| H8 | 0.0938 | 50 | 3 |
| H9 | 0.1875 | 50 | 2 |
| H10 | 0.1875 | 50 | 3 |
| H11 | 0.375 | 50 | 2 |
| H12 | 0.375 | 50 | 2 |
| H13 | 0.0469 | 50 | 4.5 |
| H14 | 0.0469 | 50 | 5 |
| H16 | 0.0938 | 50 | 5 |
| H18 | 0.1875 | 50 | 2.5 |
| H19 | 0.0938 | > 50 | > 16 |
| H21 | 0.0938 | > 50 | > 16 |
| H22 | 0.0938 | > 50 | > 16 |
| H23 | 0.0938 | > 50 | > 16 |
| H24 | 0.0938 | > 50 | > 16 |

* Calculated after 48 h due to the absence of cell growth at 24 h.

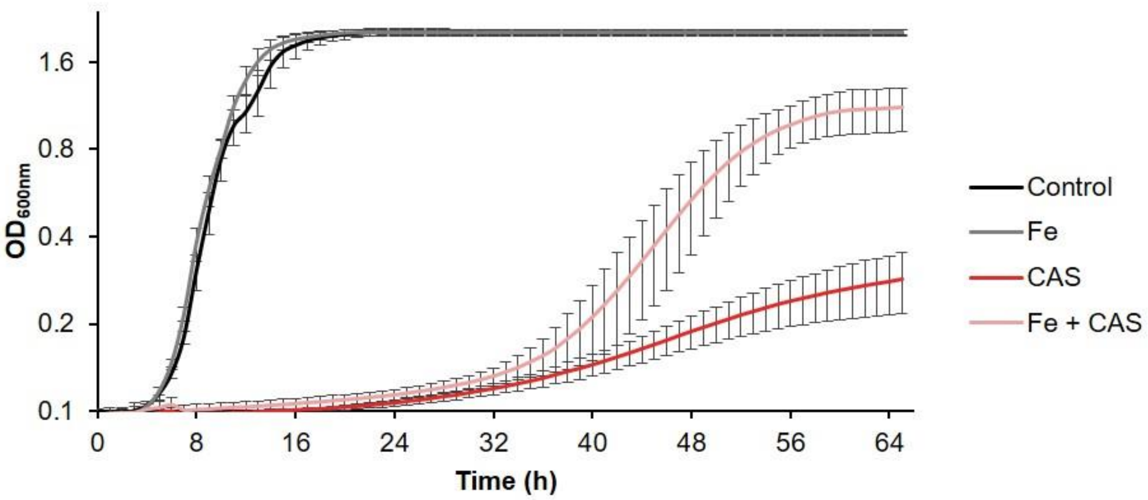


**Figure S1. Low iron concentrations reduce caspofungin toxicity in *C. albicans*.** Growth profile of *C. albicans* SC5314 in the presence of 31.25 µM $FeSO_4$ (Fe), 0.375 µg/mL caspofungin (CAS) or a combination of both (Fe + CAS). Growth was recorded for 65 h at 30 °C, by measuring $OD_{600}$ at 1 h intervals. Error bars represent mean ± standard deviation.

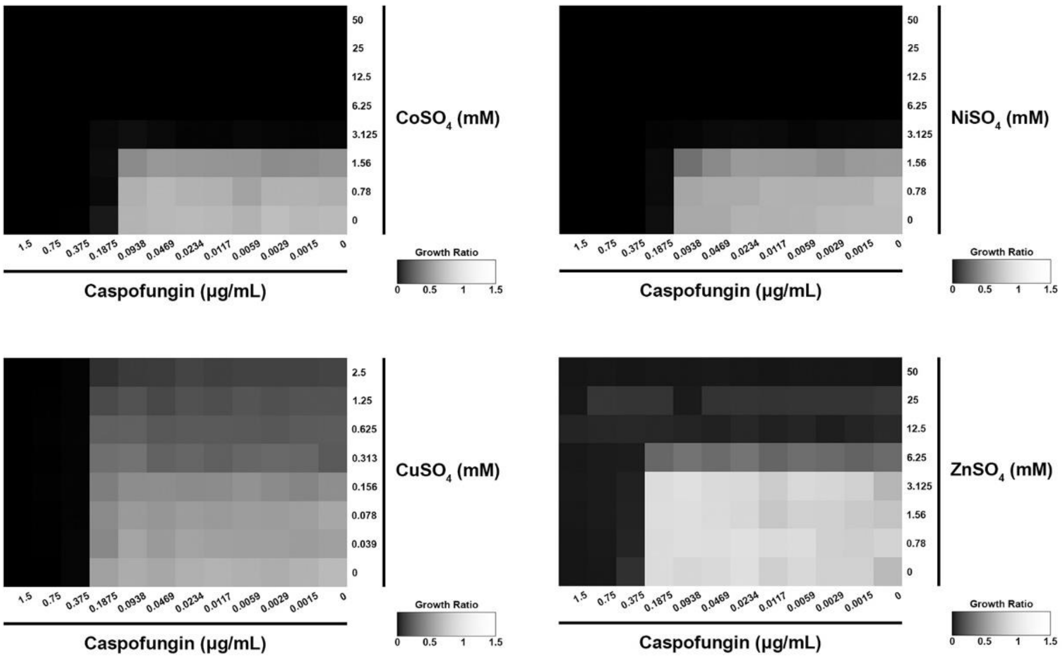


**Figure S2. The antagonistic effect between iron and caspofungin is iron-specific.** Checkerboard assays with caspofungin and either $CoSO_4$, $NiSO_4$, $CuSO_4$ or $ZnSO_4$ were performed for *C. albicans* SC5314.

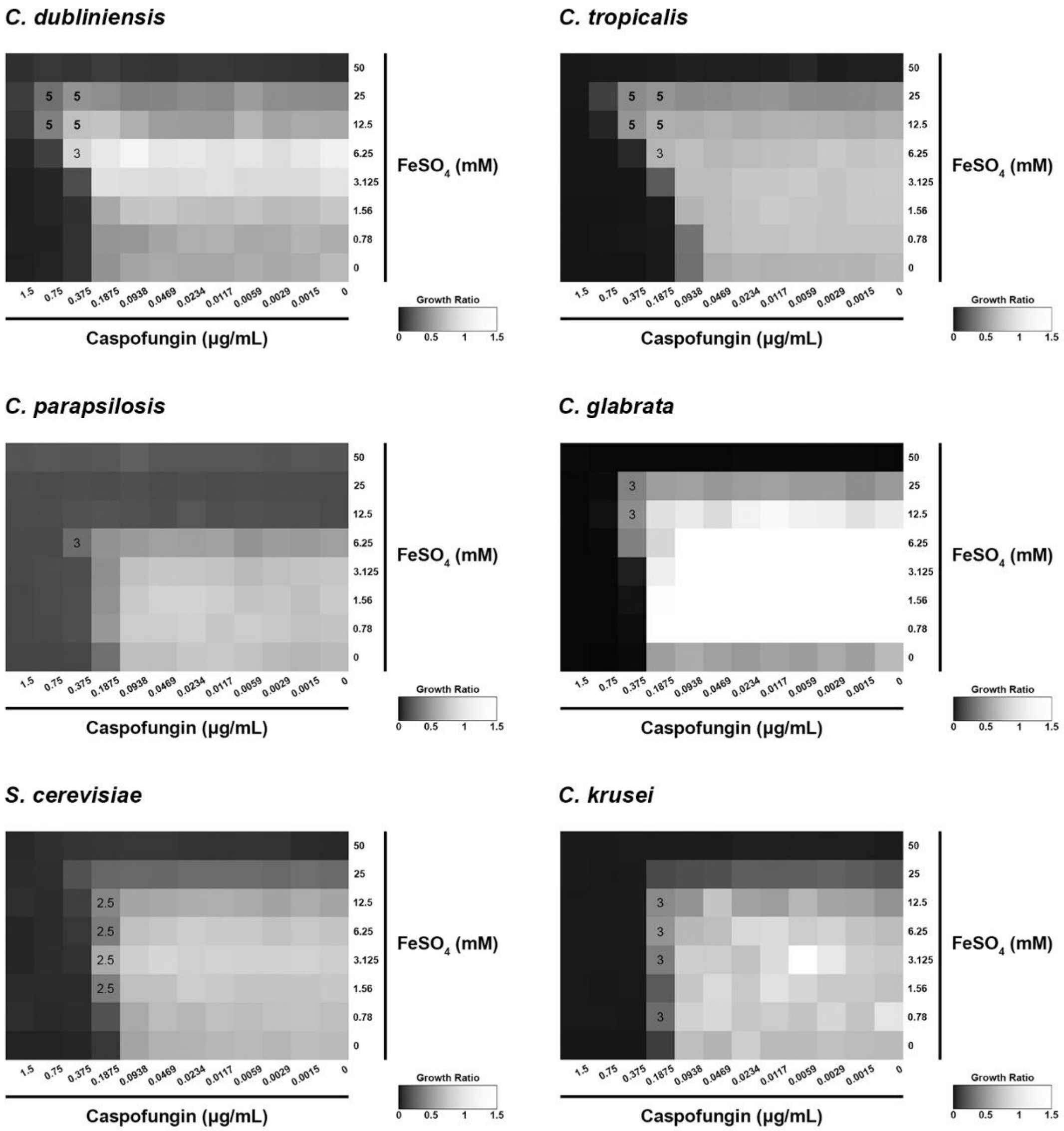


**Figure S3.** ***Candida dubliniensis*** **and** ***Candida tropicalis*** **display antagonism between iron and caspofungin.** Checkerboard assays with $FeSO_4$ and caspofungin were performed in *C. dubliniensis* MYA-577, *C. tropicalis* 7501, *C. parapsilosis* 22019, *C. glabrata* 2001, *S. cerevisiae* BY4742 and *C. krusei* 6258. The fractional inhibitory concentration index (ΣFIC) was calculated for the specified combinations of caspofungin and $FeSO_4$. Combinations demonstrating antagonism are highlighted with bold ΣFIC values.

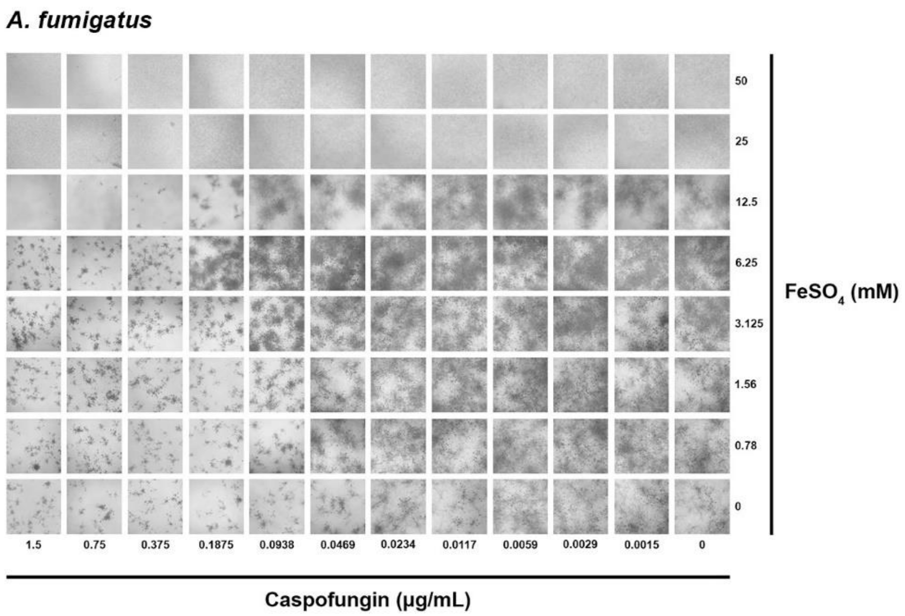


**Figure S4. *A. fumigatus* exhibits antagonism between iron and caspofungin.** Checkerboard assays with $FeSO_4$ and caspofungin were performed for *A. fumigatus*. Photos of each microplate well were taken after 24 h of incubation.

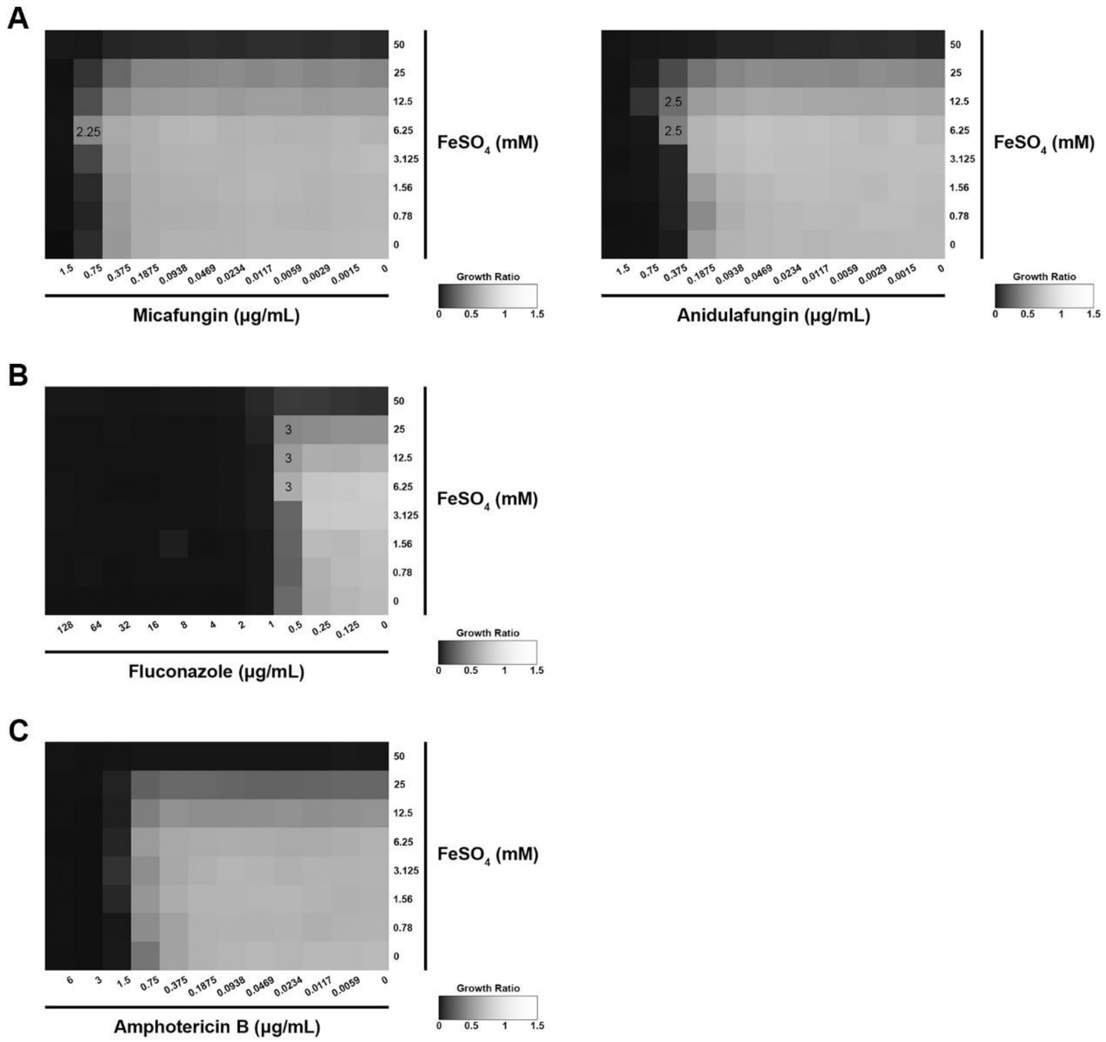


**Figure S5. The antagonistic effect between iron and caspofungin in *C. albicans* is specific to caspofungin.** Checkerboard assays with $FeSO_4$ and representatives from the (A) echinocandin class, (B) azole class, and (C) polyene class were conducted using *C. albicans* SC5314 cells. The fractional inhibitory concentration index (ΣFIC) was calculated for the specified combinations of drug and $FeSO_4$.

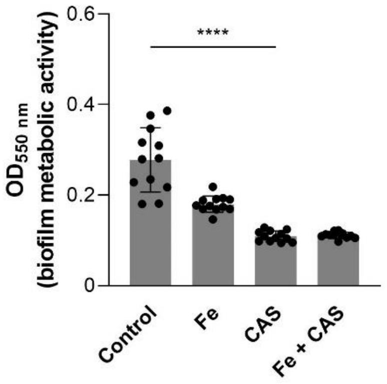


**Figure S6. Mature biofilm disruption by caspofungin is not prevented by ferric iron.** Biofilm viability of *C. albicans* was assessed for untreated biofilms (Control) or biofilms treated with 0.5 mM $FeCl_3$ (Fe), 0.1875 µg/mL caspofungin (CAS), or a combination of both compounds (Fe + CAS). Biofilm viability was quantified using the MTT colorimetric assay and by measuring $OD_{550}$. Statistical significance of differences was determined using Student’s *t*-test (**** $P < 0.0001$).

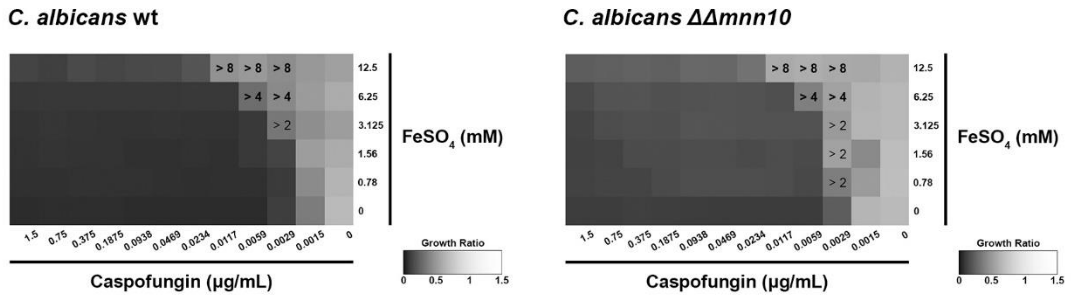


**Figure S7. Differences in mannans levels do not explain the antagonistic effect between caspofungin and iron.** Checkerboard assays with $FeSO_4$ and caspofungin were performed using *C. albicans* BWP17 (wt) and *C. albicans* SKY68 (ΔΔ*mnn10*). The fractional inhibitory concentration index (ΣFIC) was calculated for the specified combinations of caspofungin and $FeSO_4$. Numbers within the squares indicate the ΣFIC values.

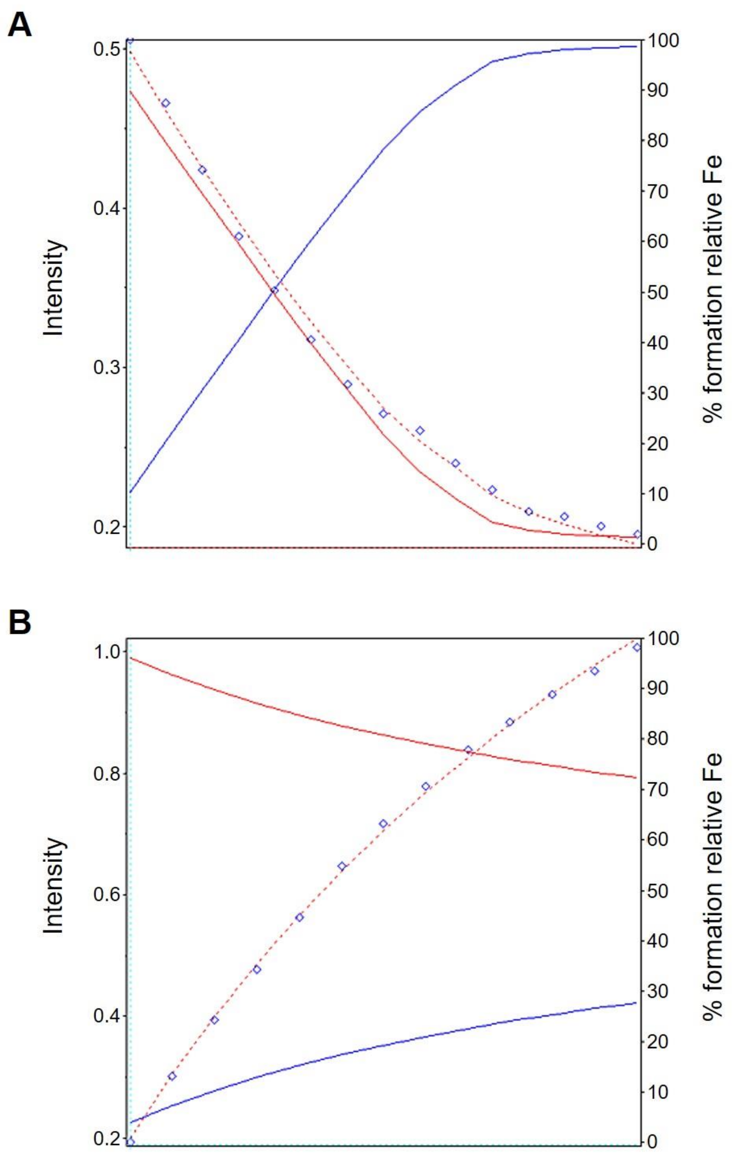


**Figure S8. Examples of the UV spectrophotometric data refinement to obtain the conditional binding constants, at a wavelength of 300 nm, for caspofungin (A) and micafungin (B).** The experimental points are marked with blue lozenges, while the red dotted line represents the data fitting. The abscissa axis represents the stepwise addition of each drug during the titration. The ordinate axes represent the absorbance intensity of the points at 300 nm (left) and the percentage of each species present in equilibrium relative to the total iron amount (right). The blue line represents the caspofungin-bound iron species, and the red line the unbound iron species.

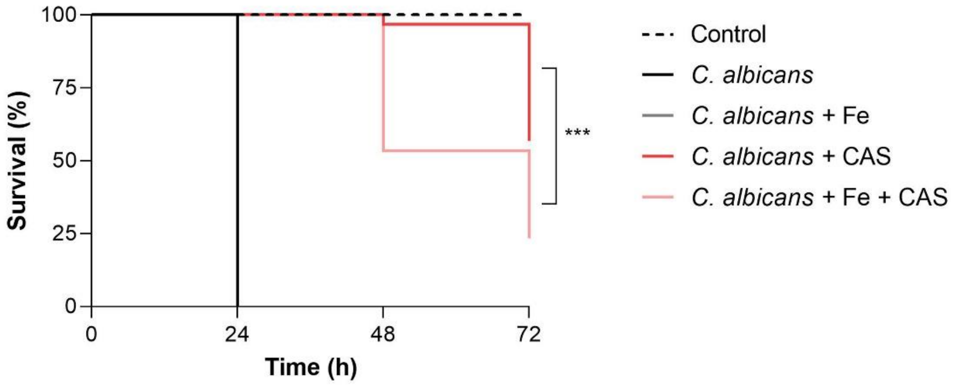


**Figure S9. Iron supplementation at concentrations as low as 100 μM impairs the *in vivo* efficacy of caspofungin against *C. albicans* in the *G. mellonella* infection model.** Survival rates of *G. mellonella* larvae injected with 0.9% NaCl (Control) or infected with *C. albicans* clinical strain H19 before (*C. albicans*) or after loading with 100 μM $FeCl_3$ (*C. albicans* + Fe), and treated with 1 mg/kg caspofungin (*C. albicans* + CAS and *C. albicans* + Fe + CAS). Statistical significance of differences in survival rates was determined using the log-rank test (*** $P < 0.001$).
